# RNA Transcriptome Mapping with GraphMap

**DOI:** 10.1101/160085

**Authors:** Krešimir Križanović, Ivan Sović, Ivan Krpelnik, Mile Šikić

## Abstract

Next generation sequencing technologies have made RNA sequencing widely accessible and applicable in many areas of research. In recent years, 3rd generation sequencing technologies have matured and are slowly replacing NGS for DNA sequencing. This paper presents a novel tool for RNA mapping guided by gene annotations. The tool is an adapted version of a previously developed DNA mapper – GraphMap, tailored for third generation sequencing data, such as those produced by Pacific Biosciences or Oxford Nanopore Technologies devices. It uses gene annotations to generate a transcriptome, uses a DNA mapping algorithm to map reads to the transcriptome, and finally transforms the mappings back to genome coordinates. Modified version of GraphMap is compared on several synthetic datasets to the state-of-the-art RNAseq mappers enabled to work with third generation sequencing data. The results show that our tool outperforms other tools in general mapping quality.

## 1 Introduction

The advent of Next Generation Sequencing (NGS) methods has popularized sequencing in various fields of research such as medicine, pharmacy, food technology and agriculture. Compared to the already established Sanger sequencing, NGS was much cheaper and much faster, while boasting similar accuracy but with significantly shorter read length. This short read length presented (and still presents) a serious obstacle for DNA data analysis, especially for larger genomes with larger and more numerous repetitive regions. The need for longer reads that could solve some of the DNA data analysis problems resulted in the development of several new sequencing technologies, jointly called “third generation sequencing technologies”.

The first of the new technologies producing longer reads was developed by Pacific Biosciences (PacBio) and named Single Molecule Real Time (SMRT) sequencing. Pacbio technology produces reads with length up to a hundred thousand base pairs, but which also have much higher error rate (∼10-15%) compared to latest NGS technologies (∼1%) [1,2].

Oxford Nanopore Technologies (ONT) presented their portable sequencer in 2014. The MinION fits in average person’s hand and is connected to a personal computer through an USB port. Read lengths it produces are similar if not higher to those of PacBio SMRT technology with the longest read reported over 200kbp long. Reported error rates for latest R9.4 chemistry are also comparable with PacBio technology: ∼10% for 2D reads and 15-20% for lower quality 1D reads (http://lab.loman.net/2016/07/30/nanopore-r9-data-release). It has been shown that even on older chemistry R7.3, bacterial genomes can be successfully assembled using solely ONT MinION reads [3,4].

Aside from DNA sequencing, NGS also enabled RNA sequencing using sequencing-by-synthesis approach. While 3^rd^ generation sequencing technologies are rapidly taking over their share of DNA sequencing market, due to the fact that read length is less important for RNA data analysis, RNA sequencing is still predominately done using NGS. However, it seems likely that at least some aspects of RNA analysis would benefit from increased read length. Some studies have shown that, even within NGS, longer reads improve mappability and transcript identification [5,6]. Long reads also enable improved “split-read” analyses so that various types of structural changes can be more easily recognized [7]. However, together with longer read length, both established third generation sequencing technologies bring a significant increase in error rate.

Currently used methods require RNA to be converted to cDNA prior to sequencing. This process has been show to introduce many biases and artefacts that interfere with proper characterization and quantitation of transcripts. Company Helicos (bankrupt since 2012) tried to address this using proprietary single molecule Direct RNA Sequencing (DRSTM) technology [8]. At the end of 2016, ONT also presented their own direct RNA sequencing technology [9] which could represent a great boost for third generation RNA sequencing.

RNA read alignment in which RNA sequencing reads are mapped either onto a genome or a transcriptome is a crucial step of most RNA analysis pipelines. If performed with high precision, this step can cover up faulty quality control and make subsequent gene and isoform abundance estimation steps much easier and much more accurate.

RNA alignment tools can be divided into two groups. De novo aligners map RNA reads to the reference genome without any prior information on gene annotations. On the other hand, guided splice aligners use known gene annotations to guide the mapping process and to calculate gene or isoform abundance. Since guided splice aligners, such as RUM [10], already use annotations in the mapping process, they cannot be used to identify new splice junctions. Some de novo splice aligners first perform initial read mapping trying to discover exon junctions, and in the second step perform guided mapping trying to improve the results. Examples of such aligners are GSNAP and GMap [11,12], MapSplice [13], TopHat2 [14] and HISAT2 [15]. BBMap [16] can be used as a DNA aligner as well as an RNA aligner. It uses short sequences called kmers to align reads directly to a genome (spanning introns) or a transcriptome.

It claims support for both ONT and PacBio reads. BBMap does not use any heuristics to find splice sites. Aside from BBMap, STAR [17] and GMap [18] (part of the same package as GSNAP) also claim support for PacBio data (but not for ONT data). Detailed instructions for using STAR and GMap with PacBio IsoSeq data are even available on PacBio GitHub pages (https://github.com/PacificBiosciences/cDNA_primer/).

Only a few RNA aligners claim support for third generation sequencing data, and only one of them specifically for ONT data. However, there are several available DNA aligners with support for long erroneous reads, such as BWA-MEM [19], Minimap [20] or GraphMap [21]. The idea that naturally occurs is, instead of using an RNA splice aligner to map RNA reads to a genome, to use a DNA aligner to map RNA reads to a transcriptome. This is the idea that we are trying to explore in this paper. In here we present an updated version of GraphMap that uses given annotations to generate a transcriptome, and then maps RNA reads to the generated transcriptome using a DNA mapping algorithm. Afterwards, the mapping results are translated back into the genome coordinates.

## 2 Methods

GraphMap is one of the first DNA mapping tools designed for long and erroneous reads, namely for PacBio and ONT data. We have recently updated it to support guided RNA spliced alignment. GraphMap takes a reference, gene annotations in GTF format and RNA reads to calculate the alignements. Due to the need for gene annotations, at this time GraphMap cannot be used for de novo spliced alignment.

### 2.1 Guided splice alignment with GraphMap

Figure 1 shows the process of RNA mapping with GraphMap. In the first step, given annotations and reference are used to generate a transcriptome. In the second step, GraphMap is used to map RNA reads to the transcriptome. Since initial alignments are calculated for the transcriptome, there is no need to consider spliced alignments and alternative gene splicing. In the third and final step, initial alignments on the transcriptome are transformed (again using given annotations and reference) back to the genome coordinates, and resulting genome alignments are output to the user. In this way, we can leverage the mapping quality of a proven DNA aligner designed for long and erroneous reads without the need for additional computation to determine exon-intron junctions.

**Figure 1.**
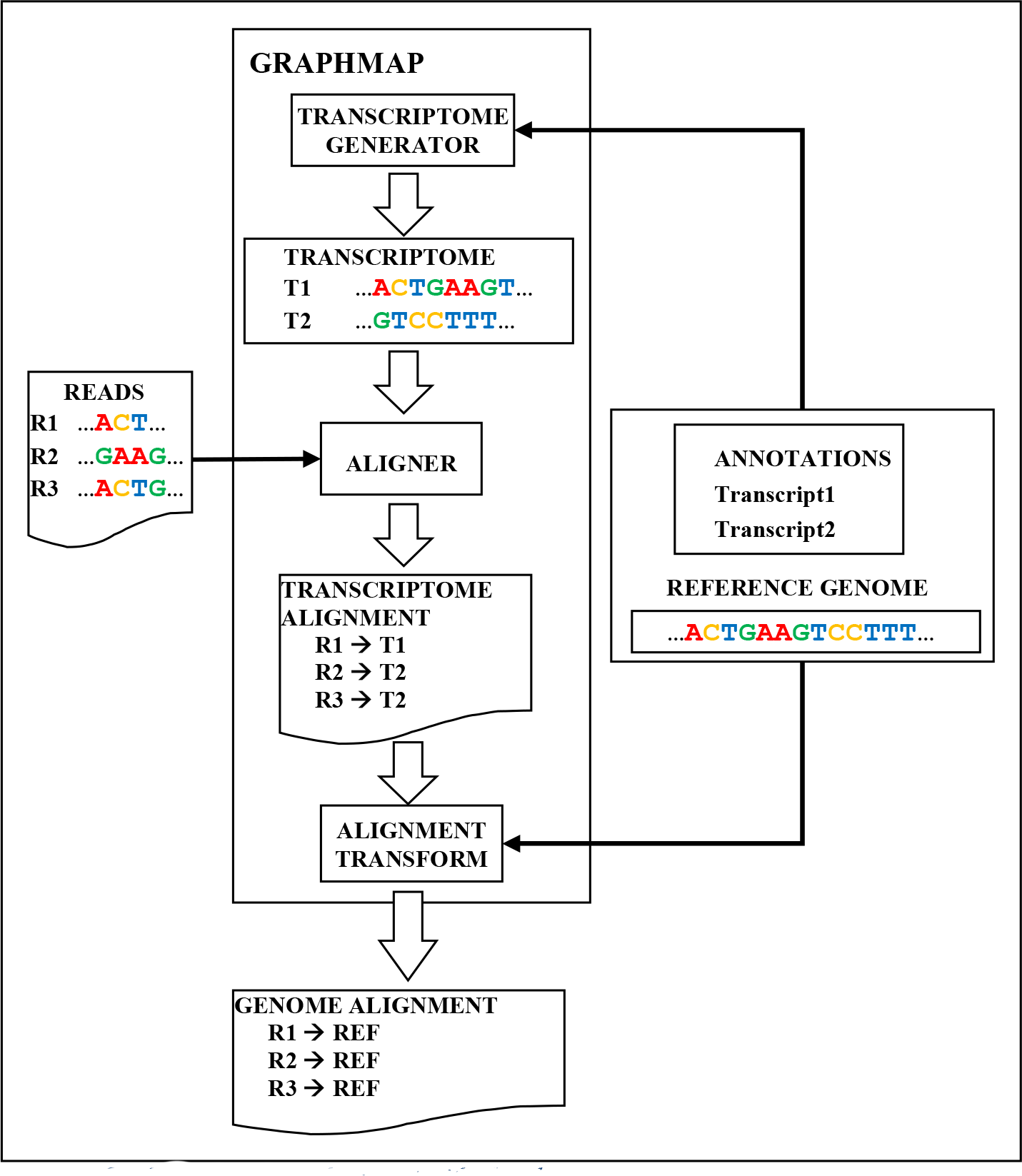
Guided RNA splice alignment with GraphMap

While DNA alignment tools have already been used for RNA mapping to transcriptome, to the best of our knowledge this is the first time the whole process has been wrapped within a single program conveniently presenting the alignments to the user in genome space.

### 2.2 Test datasets and data preparation

To be able to accurately assess the precision and quality of an aligner, read alignments must be compared to read origins. Since read origins are not reliably known for real data, we have decided to base our tests on synthetic (simulated) datasets. At the time of designing the experiments available RNA datasets obtained by third generation sequencing technologies were predominately from PacBio sequencers. Because of this, together with the fact that an appropriate simulator for ONT reads was not available, we decided to base our tests on PacBio technology.

To simulate RNA reads, we used a PacBio DNA simulator PBSIM [22]. Since PBSIM is a DNA simulator, to simulate RNA reads it was applied to a transcriptome generated from gene annotations. The number of transcripts from which the reads were generated was chosen based on real gene expression information. Annotations used to generate transcriptomes were downloaded from https://genome.ucsc.edu/. Gene expression information used to determine the number of transcripts for simulation were downloaded from http://bowtie-bio.sourceforge.net/recount/.

Test datasets were created from the following organisms:
- Saccharomyces Cerevisiae S288 (baker’s yeast)
- Drosophila Melanogaster r6 (wine fly)
- Homo Sapiens GRCh38.p7 (human)

Reference genomes for all organisms were downloaded from http://www.ncbi.nlm.nih.gov. For human genome, only chromosome 19 was used, to keep the dataset smaller and to keep the testing time within appointed time limit.

Downloaded references, gene annotations and gene expression files were used to generate three synthetic test datasets. Basic information on them is shown in (Table 1. Datasets are of different size to keep the general coverage roughly equal and because used reference genomes vary in size.

**Table 1.** Test dataset statistics

| Data set | Organism | Technology | Size | No. genes | No. reads |
| --- | --- | --- | --- | --- | --- |
| 1 | *S. Cerevisiae* | PacBio | 400 MB | 6000 | 185000 |
| 2 | *D. Melanogaster* | PacBio | 1.4 GB | 7000 | 412000 |
| 3 | Human, chr19 | PacBio | 200 MB | 1520 | 84000 |

PBSIM model for CLR reads was used for simulations, and parameters were set for PacBio ROI (Reads of Insert). Mean match rate was set to 86%, mean read length was set to 3000 base pairs and error type ratio was set to 47:38:15 (Insertions:Deletions:Mismatches).

### 2.3 Comparison with known tools

GraphMap RNA mapping was compared to 3 splice aligners that boast support for third generation sequencing data: STAR, GMap and BBMap. Since GMap and STAR are able to use information on gene annotations to improve maping results, it was provided for them.

**GraphMap:** GraphMap was downloaded from GitHub repository
https://github.com/isovic/graphmap. Version 0.5.0 was used. GraphMap was run with default parameters. Annotations were passed to GraphMap together with other parameters (option --gtf).

**STAR:** STAR version 2.5.2b was downloaded from GitHub repository: https://github.com/alexdobin/STAR. It was run with parameters suggested at *Bioinfx study: Optimizing STAR aligner for Iso Seq data* from PacBio GitHub pages (https://github.com/PacificBiosciences/cDNA_primer/wiki/Bioinfx-study:-Optimizing-STAR-aligner-for-Iso-Seq-data). Annotations were passed on to STAR during index creation (option --sjdbGTFfile). Index created using both reference and annotations was used for mapping.

**GMap:** GMap version 2016-11-07 source code was downloaded from http://research-pub.gene.com/gmap/. GMap was used with default parameters. Annotations were used to create a *map* (index) file using program iit_store, which was then added to the appropriate folder in GMap database and used during the mapping (-m parameter).

**BBMap**: BBMap version 35.92 was downloaded from https://sourceforge.net/projects/bbmap/. Script mapPacBio.sh was used, for mapping long reads. Prior to mapping reads were converted to FASTA format (originally in FASTQ format) using *samscripts* tool (https://github.com/isovic/samscripts). The mapPacBio.sh script was then run with the option `fastareadlen` set to a value appropriate for each dataset. Since BBMap is currently not able to use information on gene annotations, it was tested without it, as a de novo splice aware mapper.

Alignment results were evaluated by comparing them to MAF files containing information on read origins generated by PBSIM as a part of simulation. Evaluation was performed using a Process_pbsim_data.py script which is part of RNAseqEval package downloaded from: https://github.com/kkrizanovic/RNAseqEval. The script takes as input aligner output in SAM format, gene annotations in GTF or BED format and a folder containing files generated by PBSIM.

For each aligned read, the script finds its origin on the reference genome and compares it to the alignment calculated by the aligner. A small error in position of five bases is tolerated. The script outputs summary information on how many reads were accurately mapped to their chromosome, strand and position of origin.

Besides statistics provided by Process_pbsim_data.py script, an average match rate for each alignment file was also calculated using errorrates.py script from https://github.com/isovic/samscripts.

## 3 Results

Table 2 shows the evaluation results. Column **Mapped** shows the number of reads each aligner reported as mapped (not necessarily to the right position). Column **Match rate** shows the percentage of bases that are the same as the corresponding bases on the reference. This can be simplified as the percentage of correctly mapped bases. Column **Both ends** shows the percentage of reads for which the alignment correctly matches the beginning and the end of the read origin. Column **Hit all** shows the percentage of reads for which the alignment overlaps each exon from the read origin. Column **Hit one** shows the percentage of reads for which the alignment over-laps at least one exon from the read origin. All values (except **Match rate**) are expressed as a percentage of the initial number of reads in a dataset.

**Table 2.** Aligner evaluation results

| Dataset | Aligner | Mapped | Match rate | Both ends | Hit all | Hit one |
| --- | --- | --- | --- | --- | --- | --- |
| 1 | STAR | 49.0% | **93.7%** | 21.6% | 46.7% | 47.1% |
|  | BBMap | 91.4% | 92.5% | 48.2% | 87.0% | 88.1% |
|  | GMap | 89.1% | 92.3% | 40.7% | 84.7% | 85.7% |
|  | GraphMap | **97.4%** | 91.9% | **51.3%** | **93.5%** | **94.1%** |
| 2 | STAR | 38.2% | **94.0%** | 4.1% | 32.1% | 35.2% |
|  | BBMap | 84.5% | 89.9% | **10.5%** | 54.4% | 78.4% |
|  | GMap | 92.0% | 92.0% | 8.6% | 73.0% | 85.4% |
|  | GraphMap | **97.3%** | 91.8% | 10.1% | **82.0%** | **91.3%** |
| 3 | STAR | 37.5% | **94.3%** | 3.9% | 33.1% | 35.7% |
|  | BBMap | 64.3% | 86.2% | **10.0%** | 26.8% | 61.2% |
|  | GMap | 88.3% | 91.8% | 8.0% | 70.0% | 83.8% |
|  | GraphMap | **97.9%** | 91.8% | 9.5% | **85.7%** | **94.5%** |

Column **Both ends** represents the reads that were mapped very accurately. Column **Hit one** represents the reads that are mapped to a correct general area, but the mapping is not necessarily accurate or captures the spliced nature of reads. Column **Hit all** represents reads that are mapped to the correct general area and whose mapping captures their spliced nature, but is not necessarily very accurate.

The results clearly show that the updated version of GraphMap outperforms other RNA mapping tools in general mapping quality, having the best values in **Hit all** and **Hit one** columns for all datasets. It manages to cover all exons of the read origin for over 80% of reads on all datasets, and for over 90% on some. Average match rate is slightly lower compared to STAR on all datasets, and to GMap on datasets 1 and 2, and to BBMap on dataset 1. However, GraphMap manages to correctly map significantly more reads and lower match rate could be the result of successfully mapping lower quality reads, which other aligners were unable to map.

GraphMap also has the best or the second-best value for **Both ends** column on all datasets, however, all mapping tools map relatively small percentage of reads within the allowed error of five bases from the beginning and the end of the origin. Because of this, we believe that this value is not suitable for comparison. It is possible that 10-15% error rate is too high to achieve such highly accurate mapping.

Dataset 1 is based on baker’s yeast. It has very few spliced genes and almost no alternatively spliced genes, and is the easiest to map. All aligners except STAR obtain good results on it, and even manage to map 40-50% of the reads very accurately (column **Both Ends**). Dataset 2 is based on wine fly. It is more complex than dataset 1, with many spliced genes and some alternatively spliced ones. All mappers perform less well on it, and high accuracy mapping drops to 10% of reads and below. GraphMap and GMap still have reasonably good results, while BBMap starts to fall behind managing to hit all exons for only a little over 50% of the reads. Dataset 3 is based on chromosome 19 of human genome. All the genes are spliced and about 60% of them have alternate splicing. On this dataset GraphMap and GMap still perform reasonably well, while BBMap only manages to hit a rough area of the read origin for most of the reads. It is interesting to note that GraphMap obtains even slightly better results than on dataset 2.

Looking at the results presented in Table 2, STAR shows the lowest mapping quality of all tested mappers. It maps the least reads by a large margin, however, Match rate of the mapped reads is the highest. This could be due to STAR being able to map only the highest quality reads, which have the least errors or to STAR mapping only higher quality parts of reads, clipping lower quality starting and ending sequences. BBmap manages to get reasonable results, better than STAR, even without gene annotations. GMap shows the second best mapping results, having good results on all datasets.

## 4 Conclusion and future work

While third generation sequencing technologies are already well established for DNA sequencing, their application for RNA sequencing is still rather rare. However, with the advances in third generation sequencing technology, their more widespread use is only a matter of time. Currently, very few RNA aligners claim support for long and erroneous reads, and when put to test perform with varying success. The aim of this paper was to demonstrate another approach to mapping of third generation RNA sequencing data. The idea is to use an appropriate DNA aligner and gene annotations to map RNA reads to a transcriptome and then to transform the mapping results back to genome coordinates. While other DNA aligners could also be used for mapping RNA reads to a transcriptome, to the best of our knowledge GraphMap is the first that allows it to be done seamlessly, performing all necessary transformations internally, and the first to transform mapping back into genome coordinates, enabling their easy use for further analysis.

The research presented in this paper demonstrates that the idea is very feasible. Updated GraphMap clearly outperforms other tested splice aware aligners on all datasets. The results suggest that by implementing splice aware mapping logic into a DNA mapper which works well with third generation sequencing data could also work well for de novo RNA spliced mapping.

The results show that none of the tested RNA mapping tools can obtain high mapping accuracy, which could be due to the high error of third generation sequencing data. To further improve mapping accuracy, new approaches might be needed, such as error correcting the reads prior to the mapping.

**Funding.** This work has been supported in part by Croatian Science Foundation under the project UIP-11-2013-7353 “Algorithms for Genome Sequence Analysis“.

